# The first sheep graph-based pan-genome reveals the spectrum of structural variations and their effects on tail phenotypes

**DOI:** 10.1101/2021.12.22.472709

**Authors:** Ran Li, Mian Gong, Xinmiao Zhang, Fei Wang, Zhenyu Liu, Lei Zhang, Mengsi Xu, Yunfeng Zhang, Xuelei Dai, Zhuangbiao Zhang, Wenwen Fang, Yuta Yang, Huanhuan Zhang, Weiwei Fu, Chunna Cao, Peng Yang, Zeinab Amiri Ghanatsaman, Niloufar Jafarpour Negari, Hojjat Asadollahpour Nanaei, Xiangpeng Yue, Yuxuan Song, Xianyong Lan, Weidong Deng, Xihong Wang, Ruidong Xiang, Eveline M. Ibeagha-Awemu, Pat (J.S.) Heslop-Harrison, Johannes A. Lenstra, Shangquan Gan, Yu Jiang

**Author notes:** These authors contributed equally (Ran Li, Mian Gong, Xinmiao Zhang, Fei Wang, Zhenyu Liu, Lei Zhang).

## Abstract

Structural variations (SVs) are a major contributor to genetic diversity and phenotypic variations, but their prevalence and functions in domestic animals are largely unexplored. Here, we assembled 26 haplotype-resolved genome assemblies from 13 genetically diverse sheep using PacBio HiFi sequencing. We constructed a graph-based ovine pan-genome and discovered 142,422 biallelic insertions and deletions, 7,028 divergent alleles and 13,419 multiallelic variations. We then used a graph-based approach to genotype the biallelic SVs in 684 individuals from 45 domestic breeds and two wild species. Integration with RNA-seq data allows to identify candidate expression-associated SVs. We demonstrate a direct link of SVs and phenotypes by localizing the putative causative insertion in *HOXB13* gene responsible for the long-tail trait and identifying multiple large SVs associated with the fat-tail. Beyond generating a benchmark resource for ovine structural variants, our study highlights that animal genetic research will greatly benefit from using a pan-genome graph rather than a single reference genome.

## Introduction

Structural variations (SVs) range from 50 base pairs (bp) to over megabases (Mb) in size and are a major source of genetic variation. Compared to single-nucleotide polymorphisms (SNPs), SVs could cause larger-scale genomic perturbations in genes and regulatory regions and are therefore more likely to affect gene expression and phenotypes^1, 2^.

However, SV detection has been most challenging^3, 4^. Identification based on short reads (≤ 150 bp) has a poor sensitivity and a high false discovery rate^5^. The long reads of >10 kb by PacBio SMRT sequencing and Oxford Nanopore sequencing greatly facilitated the efficacy of SV discovery by directly spanning the breakpoints^6–11^. However, a sequencing error rate of 5-15% of the conventional long reads impede downstream analysis and is particularly problematic for analysis of highly repetitive regions. Another main restraint in efficient SV detection lines in the reference genome model. The pan-genome graph model has been shown to much superior to the linear reference assembly in representing the genome diversity and thereby facilitating analysis of large genomic variations^12–16^.

The novel PacBio circular consensus long-read sequencing (CCS) offers high fidelity (HiFi) reads of 15 to 25 kb with >99.9% base accuracy^17^, which now is revolutionizing the *de novo* assembly and SV detection. It is expected to foster the development of pan-genome applications by delivering haplotype-resolved assemblies to depict the sequence diversity of the diploid genome^18^. With the availability of population-scale genome assemblies, the pan-genomic reference models will allow to determine faithfully the structure of complex regions for reliable SV detection^7, 12, 19^.

Sheep are among the first domesticated livestock with economic and cultural importance. Previous studies have shown that SVs determine the white coat color^20^, fleece variations^21^ and polledness^22^ in sheep – characters that have been under selection for millennia. Nevertheless, a population-scale characterization of genetic variation characterized by SVs remains to be explored in sheep as in most farm animal species. In this study, we generated reference-grade *de novo* assemblies for 13 representative sheep breeds, together with two sets of phased assemblies for each individual taking advantage of the Pacbio HiFi sequencing. These datasets allow us to construct a graph-based ovine pan-genome and to explore the SV landscape with unprecedented accuracy and sensitivity. We further analyzed the SV spectrum in 684 domestic and wild sheep from short-reads sequencing and highlighted SVs as largely unexplored contributors to variations in phenotypes and gene expression.

## Results

### High-quality *de novo* assemblies of 13 representative sheep

Domestic sheep could be generally divided into at least four genetic groups including European, Asian, Middle East and Africa with a separate position of the crossbred Dorper sheep^23–25^. We selected 13 representative sheep (Fig. 1a) from seven European, four East Asian, one Middle Eastern (Kermani from Iran) and one African (White Dorper) breeds. The European breeds include meat (Suffolk, Dorset, Texel, Charollais), wool (Merino, Romney) as well as dairy (East Friesian) sheep, whereas the East Asian accessions represent the three lineages of Northern China (Ujumqin, Kazak), Qinghai-Tibetan Plateau (Tibetan) and Yunnan-Kweichow Plateau (Yunnan)^24^. A SNP analysis showed that the 13 sheep encompass 95.5% of the SNPs with minor allele frequency (MAF) >0.05 among the studied 643 domestic sheep worldwide.

**Fig. 1.**
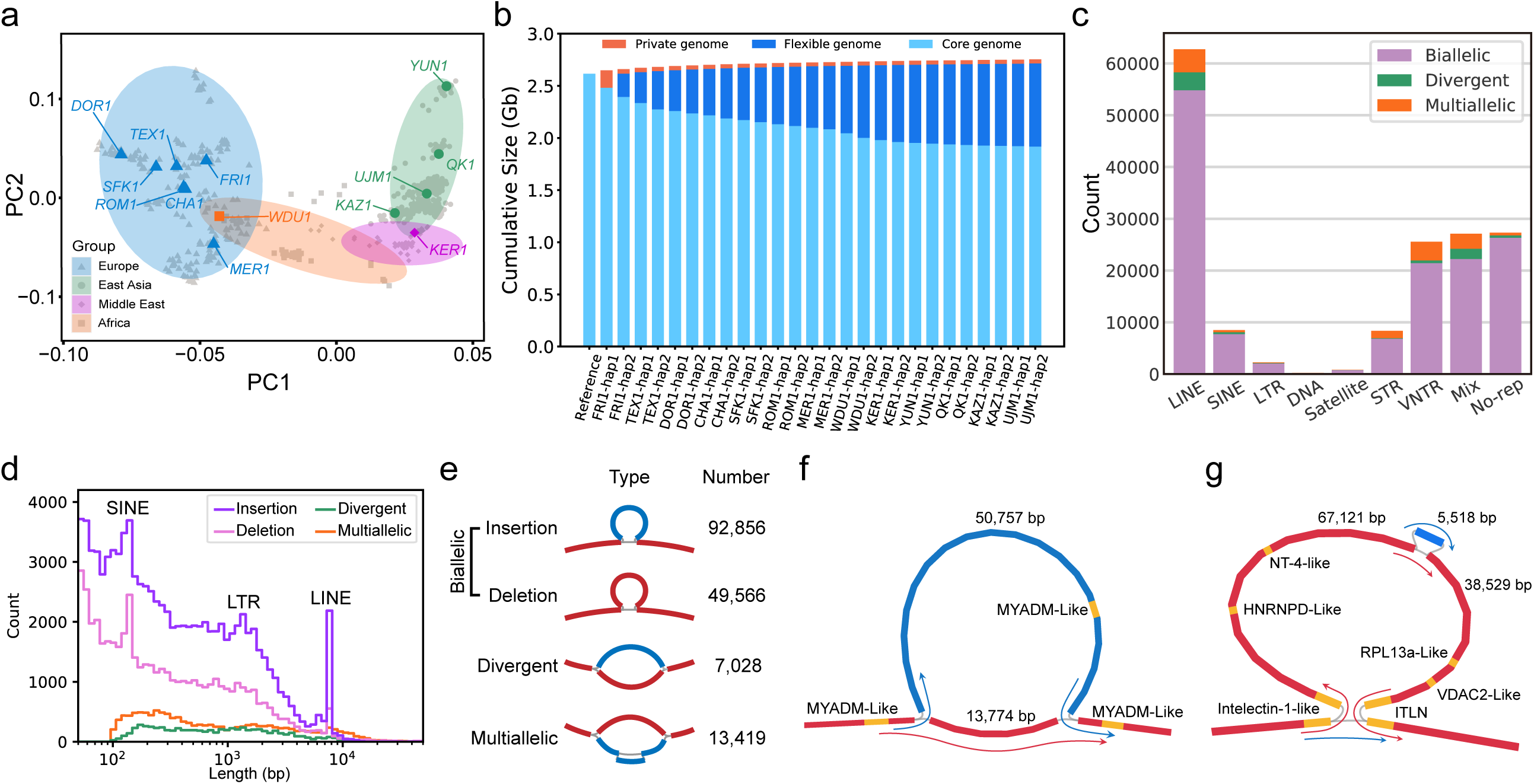
Ovine pan-genome graph construction. (a) PCA plot showing the representation of the genetic diversity of domestic sheep by the 13 individuals used for HiFi sequencing and pan-genome construction. (b) The pan-genome size changes with increase in the number of genome assemblies. (c) Repeat annotation of the structural variations stratified by SV types. (d) Length distribution of different type of SVs. (e) Diagram and number of different SV types. The red bold lines indicate the reference sequence and the blue bold lines represent the non-reference. (f) An example of a divergent allele of 50,757 bp containing an MYADM-like repeat. (g) An example of a multiallelic variation containing a large deletion of 105,650 bp which is nested with another insertion of 5,518 bp. The red lines with arrows show the reference path and the blue lines with arrows refer to the non-reference path.

The 13 sheep were sequenced using PacBio HiFi sequencing at an average coverage of 20.9×, yielding highly accurate long reads with a per-base accuracy >99.9%. We used the Hifiasm assembler^18^ for *de novo* assembly, which generated the primary contigs as well as two sets of phased contig assemblies from HiFi reads. We ultimately assembled 13 primary collapsed assemblies with an average length of ∼2.86 Gb (Table 1). The average contig number were 1582 with contig N50 lengths of 68.2 Mb. The most complete assembly (FRI1) achieved a contig N50 of 85.1 Mb, which is much higher than for the sheep reference genome (ARS-UI_Ramb_v2.0). The HiFi sequencing depth is not correlated with assembly length (*r_s_*=-0.04, Spearman rank correlation), but with contig N50 length (*r_s_* =0.93) and negatively with contig N50 number (*r_s_* =-0.67). We further ordered and positioned ∼93.6% of the contigs from each of the 13 collapsed assemblies into chromosomes based on their synteny to the reference genome. The resulting chromosome-level collapsed assemblies for the 13 representative breeds may very well serve as a reference panel for future research in sheep genetics and genomics. In addition, the haplotype phased assemblies achieved an average contig N50 length of 14.5 Mb, which captures per diploid individual almost all sequence variants and was used for the subsequent construction of the multiassembly graph.

**Table 1.** Quality assessments of the primary and partially phased assemblies for the 13 individuals. For each sheep, we show the values for the primary assembly and for both partially phased haploid assemblies.

| Breeds | Abbr. | Assembly | Assembly<br>length (Gb) | Contig<br>N50 (Mb) | Contig<br>number | Depth | BUSCO<br>(%) | Base<br>accuracy |
| --- | --- | --- | --- | --- | --- | --- | --- | --- |
| Kazak | KAZ1 | Primary | 2.87 | 73.4 | 1849 | 22.4 | 93.4 | 46.3 |
|  |  | hap1/hap2 | 2.85/2.84 | 10.7/13.9 |  | - | 92.3/91.9 | 46.6/46.4 |
| Kermani | KER1 | Primary | 2.82 | 80.3 | 1675 | 21.7 | 93.8 | 45.2 |
|  |  | hap1/hap2 | 2.77/2.75 | 15.4/15.2 |  | - | 90.7/92.2 | 45.4/45.6 |
| Dorset | DOR1 | Primary | 2.91 | 92.4 | 1293 | 25.0 | 94.3 | 45.5 |
|  |  | hap1/hap2 | 2.86/2.83 | 32.8/39.5 |  | - | 93.4/92.7 | 45.6/45.6 |
| East | FRI1 | Primary | 2.90 | 85.1 | 982 | 28.0 | 94.4 | 48.2 |
| Friesian |  | hap1/hap2 | 2.77/2.75 | 62.9/53.2 |  | - | 91.3/92.8 | 48.4/48.5 |
| Chinese | MER1 | Primary | 2.84 | 60.0 | 1773 | 18.7 | 94.2 | 43.3 |
| Merino |  | hap1/hap2 | 2.82/2.83 | 3.7/4.8 |  | - | 92.1/90.0 | 42.4/43.3 |
| Qiaoke | QK1 | Primary | 2.87 | 75.0 | 1654 | 21.5 | 94.0 | 47.1 |
|  |  | hap1/hap2 | 2.83/2.84 | 13.6/9.9 |  | - | 92.6/93.4 | 47.6/47.2 |
| Romney | ROM1 | Primary | 2.86 | 68.3 | 1552 | 19.3 | 94.4 | 43.2 |
|  |  | hap1/hap2 | 2.81/2.82 | 6.3/5.7 |  | - | 92.1/92.4 | 43.3/43.1 |
| Suffolk | SFK1 | Primary | 2.88 | 65.3 | 1513 | 18.8 | 94.3 | 43.4 |
|  |  | hap1/hap2 | 2.86/2.80 | 5.7/4.9 |  | - | 92.4/92.5 | 43.2/43.3 |
| Texel | TEX1 | Primary | 2.82 | 47.6 | 1835 | 17.9 | 94.4 | 44.6 |
|  |  | hap1/hap2 | 2.76/2.75 | 4.52/4.49 |  | - | 91.4/90.2 | 44.8/44.8 |
| White | WDU1 | Primary | 2.81 | 17.8 | 2125 | 15.4 | 93.8 | 41.7 |
| Dorper |  | hap1/hap2 | 2.82/2.77 | 1.58/1.26 |  | - | 90.2/89.8 | 41.9/41.8 |
| Ujumqin | UJM1 | Primary | 2.87 | 75.6 | 1534 | 21.3 | 94.4 | 44.4 |
|  |  | hap1/hap2 | 2.86/2.84 | 11.5/11.7 |  | - | 93.9/93.5 | 44.5/44.6 |
| Charollais | CHA1 | Primary | 2.86 | 71.5 | 1427 | 19.2 | 94.1 | 44.1 |
|  |  | hap1/hap2 | 2.82/2.80 | 6.4/5.1 |  | - | 91.7/92.1 | 44.3/44.3 |
| Yunnan | YUN1 | Primary | 2.91 | 73.9 | 1293 | 22.0 | 94.1 | 46.9 |
|  |  | hap1/hap2 | 2.89/2.85 | 18.7/13.1 |  | - | 93.6/93.4 | 46.7/46.6 |

On the basis of Benchmarking Universal Single-Copy Orthologs (BUSCO) analysis, the collapsed assemblies displayed an average completeness of 94.1% (93.8%-94.4%, Table 1). BUSCO scores for the partially phased assemblies were lower but still consistently >90% complete. To measure the quality value (QV) of the assemblies, the 13 sheep were sequenced to an average of 30× depth Illumina short-reads, which showed for all the collapsed and phased assemblies except WDU1 higher values than obtained for the sheep reference genome assembly (QV=42.5).

### Construction of sheep graph-based pan-genome and SV detection

To build a comprehensive sheep graph-based pan-genome, we used the reference genome (ARS-UI_Ramb_v2.0) as the backbone. The 26 haplotype-resolved assemblies from the 13 individuals were individually added to the graph to encompass the sequence variations more comprehensively than the primary contigs. A multigraph genome was generated using minigraph which is a state-of-the-art graph constructor that especially is suitable for discovering complex variations^26, 27^.

The resulting multiassembly graph spans 2.75 Gb and includes 167,565 nonreference nodes of 137.7 Mb. Based on the presence or absence of each node among the reference genome and the 26 haplotype-resolved assemblies, we divided the pan-genome into three categories (Fig. 1b): core (nodes present in all assemblies, 69.6%), flexible (nodes present in at least two assemblies, 29.0%) and private (nodes present in one assembly, 1.4%). The variations were most associated with LINEs followed by VNTR and mixed interspersed repeats (Fig. 1c).

The genome graph represents SVs as bubbles that are located between a single source and a single sink node coming from the reference assembly. We classified the SVs into three categories: (1) biallelic insertions and deletions; (2) divergent alleles with a non-reference node; (3) multiallelic variations with more than one non-reference path (Fig. 1d and 1e, Supplementary Data 1 and 2). The genome graph revealed 71,401 biallelic insertions and deletions, 7,028 divergent alleles and 13,419 multiallelic variations. However, despite advantage in recovering divergent alleles and multiallelc variations, the genome graph constructed by minigraph is not well adapted for the discovery of small SVs^18^ and thus we excluded those below 100 bp. As a complementary approach to detect insertions and deletions, we used pbsv (https://github.com/PacificBiosciences/pbsv) to jointly call among the 13 individuals SVs using the PacBio HiFi reads, yielding 142,279 insertions/deletions. We observed much more insertions than deletions, contrasting with results based on short-read sequencing which preferentially identifies deletions^28^. The pbsv call set covers 97.5% of the insertions and deletions found by minigraph. Therefore, we merged the pbsv call set and the large variants (>5 kb) specifically found by minigraph, resulting in 142,422 insertions and deletions, which together with the identified divergent alleles and multiallelic variations constituted a final comprehensive catalogue of SVs in the sheep genome (Fig. 1d and 1e, Supplementary Data 3).

The number of insertions/deletions rapidly decreases with length, consistent with reports in humans using long-read sequencing^6, 7, 29^. Noticeable peaks at 150 bp correspond to SINES, 1,300 bp for LTRs, as well as 7,750 bp for LINEs (Fig. 1d). The median length of divergent alleles (726 bp) and multiallelic loci (575 bp) are longer than that of biallelic variations (230 bp).

The insertion sequences were predicted to harbor 588 genes whereas the deletions potentially affect the exonic region of 1920 genes by containing exons. The nonreference sequences of the divergent alleles contained 31 genes. We revealed the full sequence of one divergent locus containing *MYADM-like* repeat (Fig. 1f), which was previously reported to be associated with mean corpuscular hemoglobin concentration traits and weight of lamb weaned but the sequence was unknown^30^. We also noticed several other large divergent alleles that likely to correspond to phenotypic variants (Supplementary Fig. 1). The non-reference alleles of the multiallelic loci were predicted to harbor 146 genes. One of the largest multiallelic region represented a deletion of 105.6 kb which is nested with an insertion of 5.5 kb (Fig. 1g). The deletion likely results in the loss of the intelectin genes, that displays extremally high expression in lung of sheep and potentially plays a role in mucosal response in allergy^31^. Additional examples of large multiallelic loci that resulted in gene gain/loss, included *HBB*, *ASIP*, *UDP-glucuronosyltransferase 2B18-like*, olfactory receptor (*LOC101115700*), etc. (Supplementary Fig. 2).

The SVs are underrepresented in untranslated region (UTR), coding sequences (CDS), and regulatory elements (H3K4me3, H3K27ac), suggesting strong negative selection that acted against them (Supplementary Fig. 3). We also reported 201 SV hotspots sites covering 18.2 Mb of the autosomes (Supplementary Fig. 4). The SVs were significantly clustered at the terminal 5 Mb region of each chromosome with 4.1-fold enrichment (*P* value<0.01, permutation test) (Supplementary Fig. 5).

### Frequency spectrum of SVs in the sheep populations

In order to better understand the frequency spectrum of the SVs within sheep, we used the graph genotyping software Paragraph^32^ to genotype the biallelic insertions and deletions in autosomes in a panel of 684 resequenced samples from 45 domestic breeds or populations and two wild ovine species. The samples included 643 domestic sheep from Europe, East Asia, Middle East and Africa, 33 Asiatic mouflons and 8 argali with an average coverage of 18.1× (Fig. 2a, see Methods for the assignment of the breeds and populations. Paragraph represent SVs as diverging paths in a graph and achieves a high rate of genotyping accuracy^33^.

**Fig. 2.**
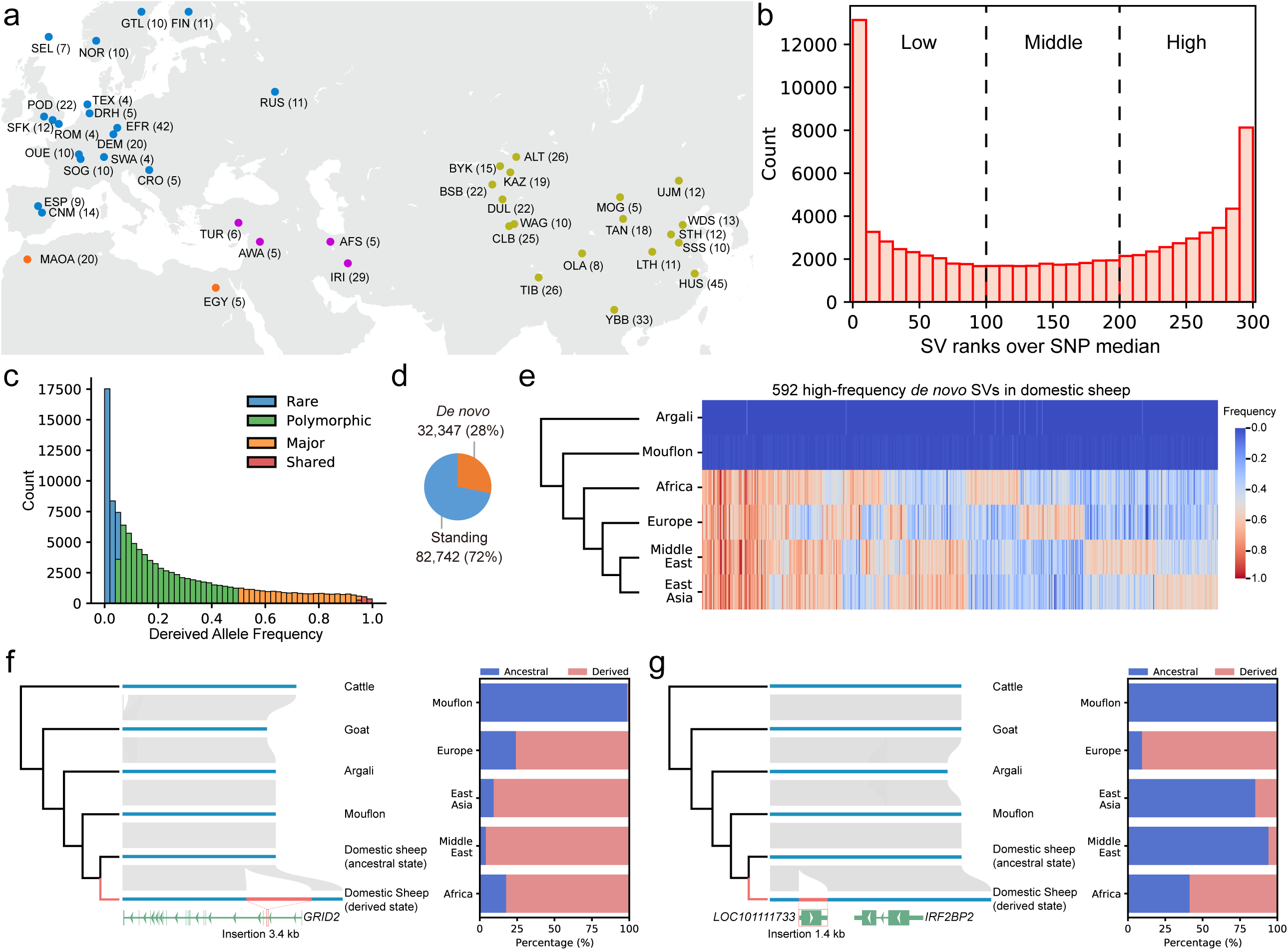
The repertoire of structural variations in sheep genome and frequency spectrum in domestic and wild sheep. (a) Geographical distribution of the 45 breeds or populations (see Supplementary Data 7 for the assignment of the breeds or populations). The Dorper sheep from South Africa and the white Suffolk from Australia are not shown on the map. (b) Histogram of the number of SV ranks (0-300) of r^2^ values that surpass the median SNP-based r^2^ value for common SVs (MAF>0.05). (c) Derived allele frequencies of the insertions and deletions. (d) The proportion of standing and *de novo* SVs. (e) High-frequency *de novo* SVs in domestic sheep. (f) A *de novo* insertion within *GRID2* occurs at high frequencies in all studied populations. (g) A *de novo* insertion of a retrogene occurs at high frequencies in European sheep.

To ensure a high-quality set of autosomal SVs for population genetics analysis, we excluded 17,330 SVs that failed to be genotyped in >90% of the samples and another 2,011 that by an excess of heterozygotes deviated from Hardy Weinberg equilibrium, leaving 116,776 (85.6%) reliably genotyped SVs for downstream analysis. The frequency of the SVs was provided in Supplementary Data 4. We also estimated the genotyping accuracy of the remaining SVs by measuring the Mendelian errors, which represent genotypes that are found in the offspring but could not be inherited form either parent. The Mendelian error rate was 3.8% on average in the 11 trios of 29 individuals (each ∼30× coverage) used in our pervious study^34^.

PCA analysis using either insertions or deletions displays similar population structure and shows a clear separation of the four genetic groups (Supplementary Fig. 6), agreeing with the clusters defined by SNPs. Therefore, the SV catalogue reproduces the genetic diversity of the worldwide sheep population.

The level of linkage disequilibrium (LD) between SVs and SNPs could reflect the extent of SVs representing independent and undiscovered genetic variations relative to previous evolutionary studies. We estimated how frequently the SVs were linked to the flanking SNPs (Fig. 2b). Around 39.8% SVs link strongly with surrounding SNPs (rank>200), the characteristic of which could be detected by the surrounding linked SNPs. It is worth noting that there are still 39.2% SVs display low levels of LD with nearby SNPs (rank<100), implying variants that are poorly tagged by easily genotyped SNP markers.

In order to infer the allele state (ancestral or derived) for each SV, we additionally genotyped one snow sheep (*Ovis nivicola*) and one bighorn sheep (*Ovis canadensis*) and, together with one argali (*Ovis ammon*) as outgroup, and identified the state for 115,089 SVs and a variable allele frequency of derived alleles (DAF) in domestic sheep (Fig. 2c). About 28% of the derived alleles occurs only in domestic sheep (Fig. 2d) and may have reached a high frequency by domestication and improvement. The proportion of *de novo* SVs may be overestimated due to the relatively limited number of wild sheep but this does not explain a high geographic differentiation of 592 high-frequency *de novo* SVs (DAF ≥ 0.5 in at least one geographic group (Fig. 2e). Those SVs were found to be enriched in KEGG pathways of Ras signaling and long-term depression (adjusted *P* value<0.05). For example, we found in all regions a *de novo* 3.4 kb insertion in the first intron of *GRID2* (Fig. 2f), which plays a key role in synaptogenesis and neurodevelopment^35^. A 1.4 kb *de novo* insertion prevalent in European sheep was found at the 3’ UTR of *IRF2BP2* (Fig. 2g), which was reported as the causative mutation for the wool coat^21^.

### Population-stratified SVs and their potential effect on gene expression

After initial domestication at the Fertile Crescent, domestic sheep dispersed worldwide and adapted to a wide range of natural and artificial selection. In this context, we detect putative selection signals by dissecting population-stratified SVs among the selected 33 out of the 45 breeds or populations to ensure a similar number of breeds or populations for European and East Asian sheep. For the differentiation between these groups, we used a modified calculation of global *F*_ST_ (see Methods). Because this strategy tends to discover common selection signals present in multiple breeds, the population-stratified SVs differentiating Europe and East Asian breeds were preferentially detected since they are in our panel the predominant breeds.

We identified 865 population-stratified SVs with the top 1% global *F*_ST_ values (Supplementary Data 5), 351 of which intersect with genic regions and 324 are within 100 kb of gene flanking region (Fig. 3a). The majority of them (698, 80.7%) were confirmed by the Ohana approach^36^ with which we detected selection signals in multiple populations by varying the number of ancestral admixture components. The related genes are associated with diverse phenotypic and production traits, including wool type (*IRF2BP2*, *FGF7*)^21, 37^, horn type (*RXFP2*)^38^, fat deposition (*BMP2*, *PDGFD*, *PDGFA*)^23, 39^, and coat color (*KIT*)^40^, etc. Notably, the highest stratified SV (*F*_ST_=0.78) is the 1.4 kb insertion downstream of *IRFBP2* (Fig. 2g), differentiating European and Asian breeds. Functional annotation of the *top F*_ST_ signals reveals enrichment in several KEGG pathways, such as those involved in signaling pathways (Rap1, MAPK) and parasitic disease (African trypanosomiasis, amoebiasis) (Supplementary Fig 7). SVs may impact the gene expression of neighboring genes via the cis-regulatory elements^10, 41, 42^. To explore SV-associated expression changes, we used 112 RNA-seq data from 8 tissues of 14 hybrids of Texel×Kazak sheep as well as WGS data of the same sheep^34, 43^ (see Methods). This unique combination of data enabled us to identify the candidate expression-associated SVs by allelic-specific expression (ASE) mapping. By ASE mapping, we found 1,141 heterozygous SVs that would putatively act as cis-regulatory elements and thus cause allelic-specific expression in associated genes (Fig. 3b and 3c, Supplementary Data 6). These expression-associated SVs were significantly enriched in generic regions (*P* value<2.2e-16, permutation test), and 34 overlapped with population-stratified SVs which is greater than random chances (*P* value<1.1e-10, Chi-Square test). We additionally performed SV-eQTL mapping and found 260 SVs with genotypes that correlate with expression levels of a nearby gene (Supplementary Data 7).

**Fig. 3.**
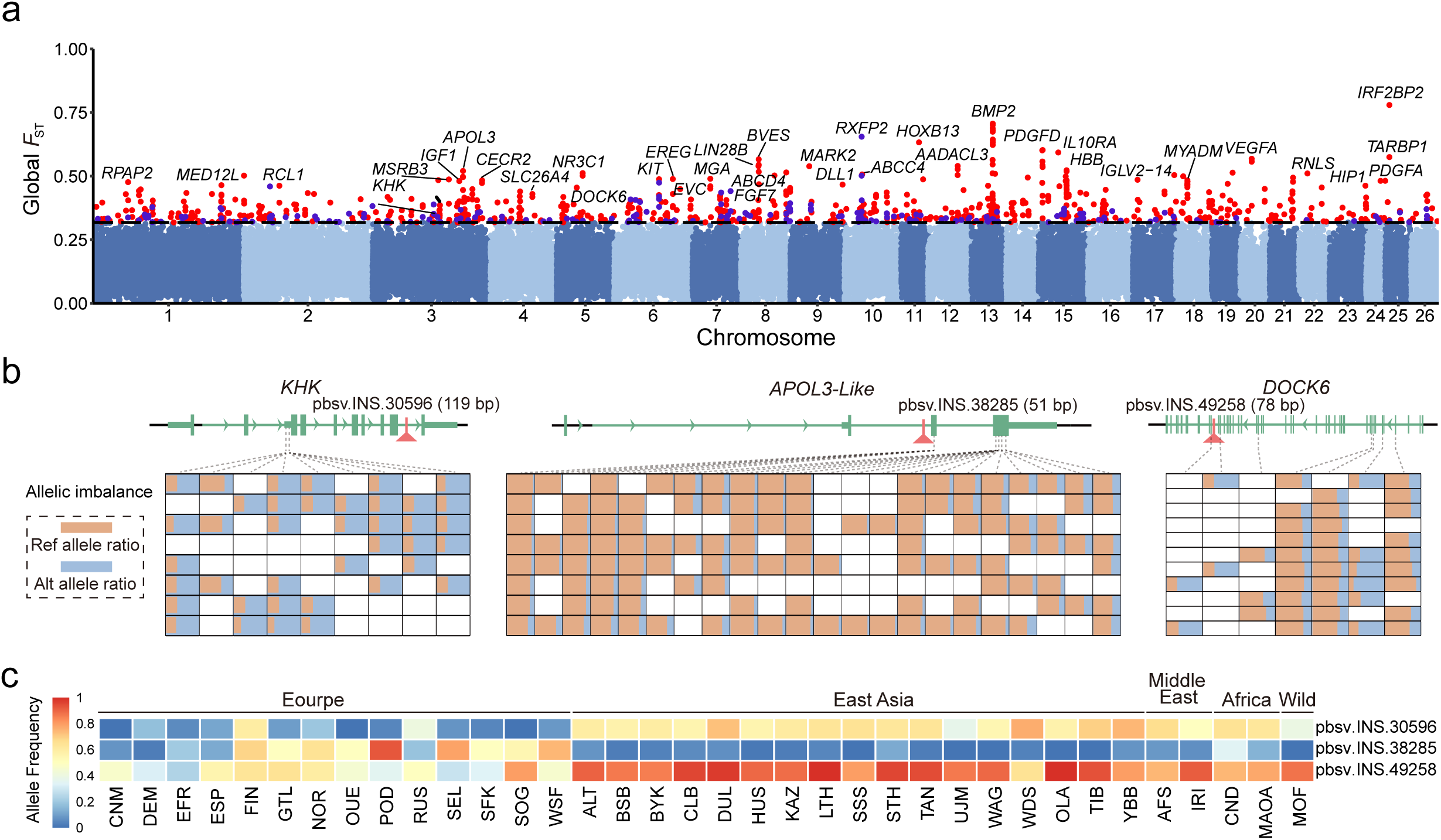
Population-stratified SVs and their effects on gene expression. (a) Genome-wide distribution of global *F*_ST_ which is measured by the average value of the top 50% *F*_ST_ values for each SV across 33 sheep breeds or populations. The red dots represent population-stratified SVs that were also found by Ohana approach. (b) Examples of expression-associated SVs identified by allelic specific expression (ASE) mapping. Each column represents one ASE SNP in exons and each row represents one RNA-seq sample. (c) Allele distributions of the three expression-associated SVs in diverse breeds or populations.

### SVs as strong candidates affecting sheep tail morphology

The tail morphology of sheep is a key genetic trait that is related to domestication, adaptation, and animal welfare^44^. The wild ancestors (Asiatic mouflon) have a short thin tail whereas four domestic tail types include in addition to the primitive type the long thin-tailed, the short fat-tailed (including fat-rumped) and the long fat-tailed types. Although several genes have been found to be associated with these traits^23, 45, 46^, the underlying causative mutations are still unknown. However, the long and fat tails are derived traits, indicating that the causative mutations are likely to be *de novo* variants.

To identify SVs associated with tail length, we compared the long and short tailed breeds of (1) the Asian long fat-tailed breeds versus Asian short fat-tailed breeds from Asian breeds and (2) the European long thin-tailed breeds versus the North European short thin-tailed breeds, respectively. Notably, in both independent comparisons the most differentiated SVs point to an insertion of 169 bp (chr11:37525005) close to the 5’ UTR of *HOXB13* (Fig. 4a). This resides in a specific haplotype exclusively found in long tailed sheep, where the insertion is prevalent in long-tailed breeds and rare in short-tailed breeds (Fig. 4b and 4c). *HOXB13* is mainly expressed in the embryo and colon of sheep and cattle^47^. We found the expression of *HOXB13* in colons of the short-tailed Texel×long-tailed Scottish Blackface individuals presumably carrying at least one copy of the insertion^48^, and in colons of individuals with short tail presumably without the insertion (Texel sheep and small-tail sheep)^49, 50^. The insertion was only expressed in the colons of the Texel×Scottish Blackface but not in Texel sheep and small-tail sheep (Fig. 4d and Supplementary Fig. 8).

**Fig. 4.**
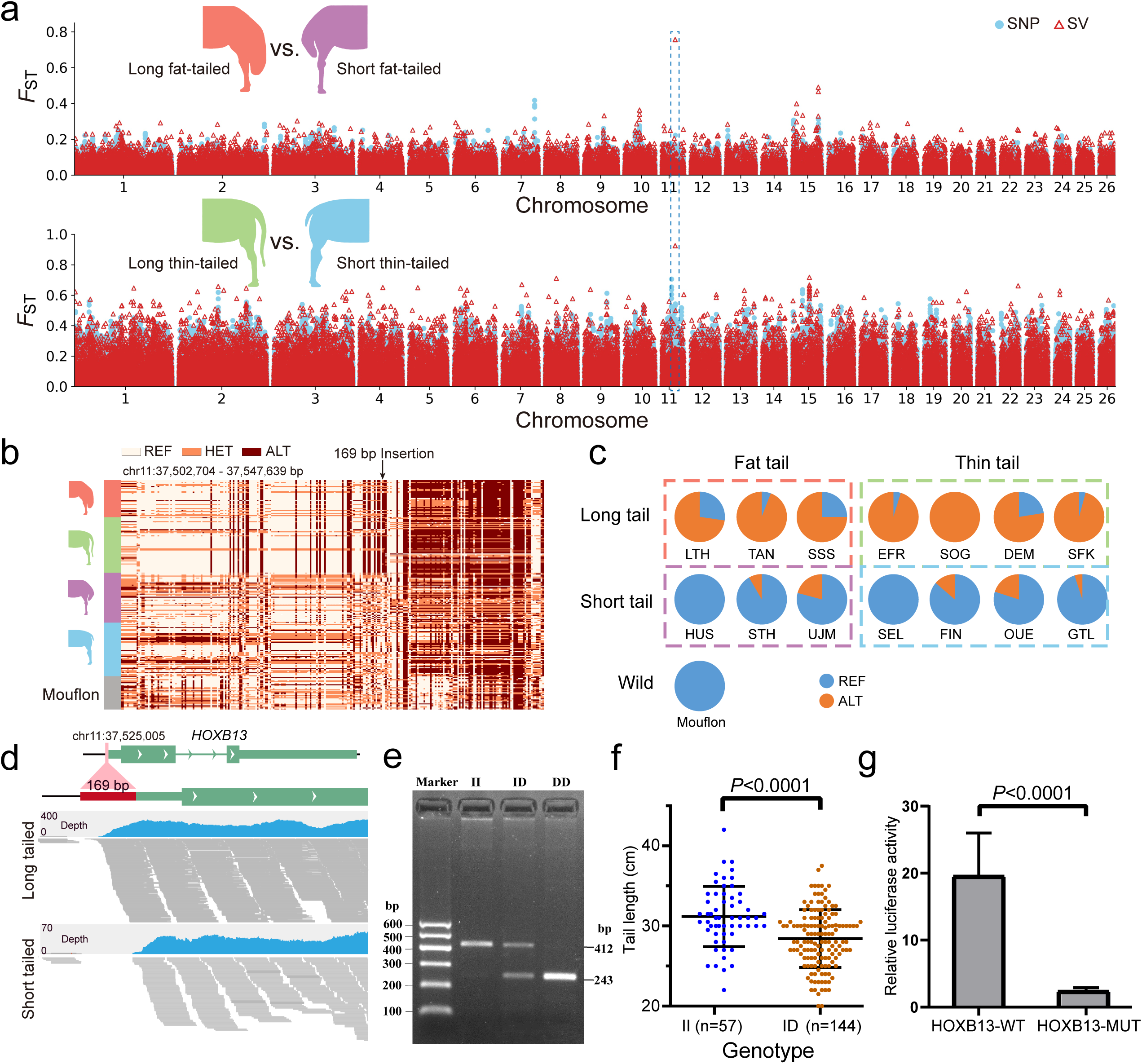
Selective SVs associated with the long tail trait. (a) The genome-wide Fixation index (*F*_ST_) of long fat-tailed versus short fat-tailed Asian sheep breeds and long thin-tailed versus short thin-tailed European sheep breeds. (b) The selective haplotype in long-tailed sheep. The 169 bp-insertion is located upstream of the 5’ UTR region of *HOXB13*. The haplotypes include the genotypes of the insertion and surrounding SNPs. (c) The frequency of the insertion in wild sheep and domestic sheep. (d) RNA-seq data shows that the expression of the insertion. The top panel corresponds to the expression in the colon of Texel × long-tailed Scottish Blackface sheep (ERS2107126); the bottom panel corresponds to the expression in colon of one small-tail sheep (SRR5122597) without the insertion. (e) The electrophoresis diagrams of PCR products for the three genotypes (homologous insertion: II, heterozygote: ID, homologous deletion: DD). (f) The carriers and non-carriers of the insertion differ in tail length. (g) *HOXB13* promoter activity is modulated by the 169 bp insertion. Luciferase activity in HEK293T cells transfected with recombinant plasmids containing the insertion or not. Experiments were repeated at least five times. The data are presented as mean ± standard error of the normalized luciferase activity.

In order to verify the effect of this insertion on tail length, we evaluated the association between the insertion and tail length in a population of 201 sheep of the same age (nine months old). The three genotypes including homologous insertion (II), heterozygote (ID) and homologous deletion (DD) were directly distinguished through PCR amplification and agarose gel electrophoresis (Fig. 4e). Despite lacking the DD individuals in the studied population (see Methods), the tail length of the homologues (II) was found to be significantly longer than that of the heterozygotes (ID) by 2.77 cm on average (Mann-Whiney U test, *P* value<0.0001) (Fig. 4f). To validate the function of the insertion, we performed luciferase reporter assays. Relative to the wildtype (WT), the mutant with the insertion decreased luciferase activity by 10-fold (Fig. 4g), indicating that the insertion reduces the transcriptional level of *HOXB13*. The gene has been previously identified as a repressor of caudal vertebra development^51, 52^. Therefore, the insertion is likely to increase the tail length of sheep by depressing the expression of *HOXB13*.

To identify candidate SVs related to the fat tail, we compared the genomes of eight fat-tail breeds (SFK, KAZ, HUS, UJM, LTH, TAN, BSB, SSS) and ten thin-tail breeds (FIN, DEM, EFR, GTL, SEL, SFK, SOG, POD, TIB, YBB). We applied the population branch statistic (PBS) to identify positive selection in fat-tail sheep using Asiatic mouflon as an outgroup. The two most differentiated regions are identified by either SV or SNPs (Fig. 5a and 5b), one located at the intergenic region between *BMP2* and *HAO1* (hereafter referred to as IBH region) and the other in *PDGFD*. Notably, the most stratified SVs in these two regions are among the top selective variants indicated by both SVs and SNPs. The most differentiated SV cluster are located at IBH region which has also reported as a candidate selective region affecting fat tail trait^53, 54^. Notably, six highly differentiated SVs including one *de novo* SV were identified in the IBH selective region (Fig. 5c), of which the largest one is a 7728 bp insertion. *PDGFD* has been recognized as the most plausible candidate gene for the fat-tail phenotype in recent studies^23, 55^. We found three highly differentiated SVs residing in the selective region of *PDGFD*, of which the largest event is a 867 bp insertion. The three selective variants are all domestic *de novo* SVs. Therefore, our results suggest that both regions are associated with the fat-tail phenotype and the SVs identified within these regions are the most plausible candidate causative mutations.

**Fig. 5.**
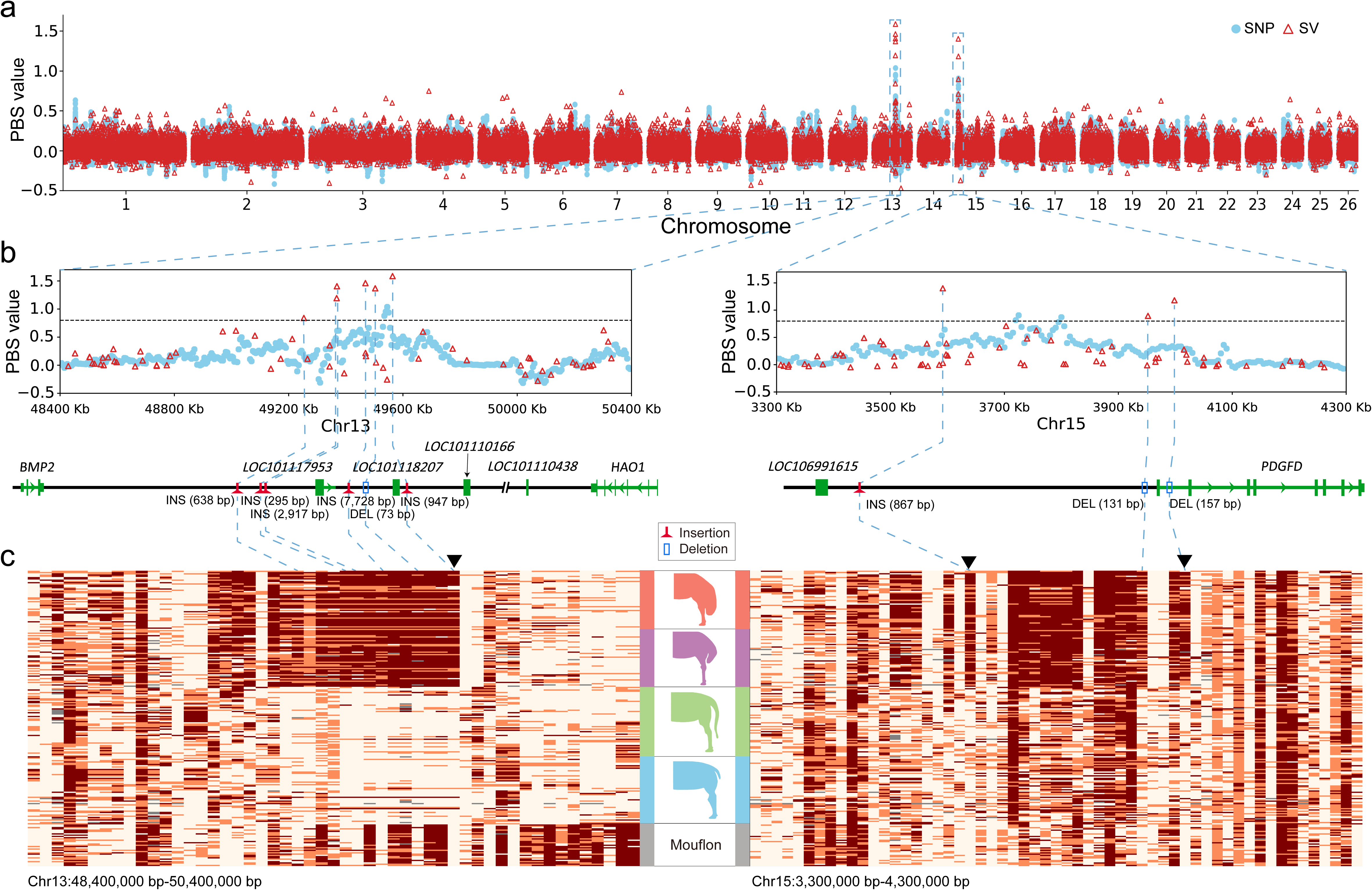
Selective SVs associated with the fat-tail trait. (a) Population branch statistic (PBS) values across the whole genome by comparing fat-tailed sheep to thin-tailed sheep using the mouflon sheep as an outgroup. (b) The two most differentiated regions between fat-tailed sheep and thin-tailed sheep. The left panel shows the intergenic region between *BMP2* and *HAO1* while the right panel corresponds to the region surrounding *PDGFD*. SVs with PBS value >0.8 are highlighted by blue dotted lines. (c) The haplotypes of mouflons and domestic sheep for the two most selective regions. Each column represents one SV and each row represent one individual. The black reverted triangles represent *de novo* SVs.

## Discussion

The genome-wide analysis of genomic structural variations in diverse breeds is critical to understanding the full repertoire of genetic diversity and pinpointing their associations with phenotypic traits. However, previous resequencing studies mainly rely on mapping short reads to a linear reference genome, precluding the discovery of regions with pronounced structural variations^4^. The graph-based pan-genome model largely reshapes the methodology to assess the whole landscape of sequence diversity, and is likely to transform the genomic research field. To build a high-quality ovine pan-genome, we utilized the PacBio HiFi sequencing to generate a panel of reference-level assemblies and 26 phased haplotypes for 13 representative sheep breeds, providing a solid foundation for in-depth functional genomic studies. The long and accurate HiFi reads guarantee the high-standard of our pan-genome through generation of high-quality *de novo* assemblies, preservation of the two haplotypes and precise assignment of SV’ breakpoints. The development of this ovine pan-genome will contribute to shifting in animal genomics research from a linear reference genome to a pan-genomic model comprising multiple reference-quality representative genomes.

The application of graph-based pan-genome has its clear opportunities and obstacles. The pan-genomic approaches are in practice not so straightforward as the linear reference genome and the methodologies and implementations are still in active development^12, 14, 56, 57^. For example, minigraph is one of the most promising graph constructors, but has limitations in performing base alignment, generating relatively high false positives for small SVs^26^. Nevertheless, a shift to the graph-based pan-genomes model will enhance the opportunities to systematically investigate previously inaccessible genomic variations. In our study, we found that the minigraph performed very well in discovering divergent alleles and multiallelic variations, which accounted for 27.4% of the total variations in length but are previously neglected. Combining the graph-based pan-genome with read-based calling yielded a reference panel of high-quality SVs for downstream functional genomic studies.

We also demonstrated how these pan-genome datasets can be utilized to link SVs to phenotypes by genotyping and assessing the SV landscape in a large population from short-read sequencing. We genotype SVs using the graph-based Paragraph, which enables efficient encoding of structural variation into the pan-genome^32^. The population-scale study allows us to trace their responses to natural and artificial selection. Taking the tail morphology traits as an example, we identified a plausible causative mutation in *HOXB13* for the long-tail trait, which illustrates the key roles of SVs underlying phenotypic changes. We further reported segregating SVs in fat-tailed breeds. The observation that *de novo* SVs showed higher or comparative selection signals than surrounding SNPs warrants further investigation.

In conclusion, the pan-genome representation with an influx of high-quality assemblies will revolutionize our understanding of SV prevalence and their biological significance, giving a new dimension to the nature and extent of genomic variation, beyond SNPs, impacting traits.

## Methods

### Sample collection and PacBio sequencing

Samples were selected to for an optimal representation of the global sheep diversities. The sampling information were provided in Supplementary Table 1. High-quality genomic DNA (gDNA) was extracted from fresh blood samples as previously described^58^ and assessed for purity and quantity using Nanodrop 1000 (Thermo Fisher Scientific, CA, USA) and Qubit (Thermo Fisher Scientific, CA, USA) assays. Library with an average insert size of ∼15 kb was generated using the SMRTbell Express Template Prep Kit 2.0 (Pacific Biosciences, CA, USA) and fractionated on the SageELF (Sage Science, Beverly, MA, USA) into narrow library fractions. Libraries were then sequenced on 2-3 SMRT Cells 8M on a Sequel II instrument (Pacific Biosciences, CA, USA) using 30-hour movie times at Annoroad Gene Technology Co., Ltd (Beijing, China). Raw data was processed using the CCS algorithm (version 6.0.0, parameters: --minPasses 3 -- minPredictedAccuracy 0.99 –maxLength 21000) to generate highly accurate HiFi reads.

### *De novo* assembly using HiFi reads and quality assessment

Hifiasm v0.15.3-r339 was used to generate the assembly from HiFi CCS reads using default parameters^18^. Hifiasm yields one primary contig assembly and two sets of partially phased contig assemblies. The primary contig assembly for each breed was scaffolded to chromosome level using RagTag v2.0.1 with the reference sheep genome assembly^59^.

The genome completeness was assessed using the BUSCO program (v.3.0.2) containing the Mammalia odb9 set of 4,104 genes^60^. The single plus duplicated complete BUSCO gene counts are reported. The base accuracy was measured by assembly quality value (QV) using yak (https://github.com/lh3/yak), which compares the 31-kmers found in short-reads and the assembly sequences.

### Pan-genome construction

The pan-genome construction was performed using a previously reported workflow^15^ with minor modifications. Minigraph v0.15-r426 with option -xggs was used to generate the graph pan-genome^26^. The sheep reference genome (ARS-UI_Ramb_v2.0, GCF_016772045.1) was used as the backbone of the multiassembly graph and the 26 partially phased contig assemblies were added successively. Only autosomal sequences were considered in the multigraph genome, as the X and Y chromosomes of males were underrepresented.

### Identification of non-reference divergent alleles and multiallelic alleles from the pan-genome graph

The graph generated by minigraph is composed of chains of bubbles with the reference as the backbone. We first used gfatools v0.5-r234 which is a bubble popping algorithm to extract bubbles from the graph (https://github.com/lh3/gfatools). Each bubble represents a structural variation, encompassing the start and end nodes of reference sequences as well as path traversing the start and end nodes.

Depending on the number of paths in a bubble, the structural variations were classified into three categories: (1) biallelic variations (insertions/deletions): either the reference path is longer (deletions) or the non-reference path is longer (insertions). The node in the shorter path is required to be below 100 bp. (2) Divergent alleles: the reference nodes and the non-reference nodes were both longer than 100 bp. (3) Multiallelic alleles: one reference path and more than one nonreference path were present. Those nonreference paths below 100 bp were excluded.

### Genome annotation of non-reference sequences

We used RepeatMasker v4.0.5 (http://www.repeatmasker.org) to softmask the non-reference sequences. Then we performed an ab initio gene structure prediction using Augustus v3.3.3^61^ and searched the protein sequence predicted by Augustus against the local protein database using DIAMOND^62^ blastp (parameters: --more-sensitive -- evalue 1e-10 --max-target-seqs 1) and kept hits with minimum coverage of 70% and identity of 80%. The local protein database was built using DIAMOND^62^ makedb with RefSeq protein sequences from sheep, goat and cattle.

### Annotation of repeats

Interspersed repeats and low complexity DNA sequences were identified using RepeatMasker v4.0.7 (http://www.repeatmasker.org) with a combined repeat database including Dfam v.20170127 and RepBase v20170127 with parameters: -species Ruminantia -xsmall -s -no_is -cutoff 255 -frag 20000 -gff. Tandem repeats were identified using Tandem Repeat Finder v4.09^63^. The identified tandem repeats as well as those low complexity sequences identified by RepeatMasker were merged as low-complexity regions (LCR). Furthermore, LCR with unit motif lengths ≥7 bp were classified as VNTR and those with unit motif lengths ≤6 bp were classified as STR.

### Structural variant calling

We integrated the results from graph-based pan-genome analysis and alignment-based SV calling to generated high-confidence SV calls. For pan-genome analysis, the SVs were generated from our multiassebly graph genome using gfatools v0.5-r234 as described above. For alignment-based SV calling, the SVs were detected using pbsv tools v2.6.2 (https://github.com/PacificBiosciences/pbsv). The CCS reads were aligned to the reference genome using pbmm2 to generate the alignment BAM file.

Then the pbsv discover module was used to identify signatures of structural variation within each aligned BAM. The tandem repeat annotation file was provided to increase the sensitivity and recall of this step. Finally, the signatures of all the samples were fed to the pbsv call module to jointly call SV from structural variant signatures using the –ccs mode.

### Mendelian error rate estimation

A Mendelian inheritance error represents a genotype in the offspring that cannot be inherited from either of their paraments. The whole genome sequencing data of 11 trios were from our recently study for allelic specific expression analysis^34^. For each trio, the Mendelian error rate is the proportion of SVs from the children that deviate from Mendelian inherence.

### SV hotspot analysis

SV hotspot regions were determined using primatR package “hotspotter” section (bw=200000, num.trial=1000, pval<1e-8)^64^. SVs were significantly clustered at the terminal 5 Mb region of each chromosome (*P* value<0.01, permutation test, random shuffling of 1,000 times). The random shuffling of SV intervals was performed using BEDTools v2.29.0 with the shuffle command^65^.

### Enrichment of SVs in the genome

The H3K4me3 and H3K27ac peak regions were retrieved from our Ruminant genome database. We only used insertions and deletions for enrichment. For insertions, the end position is the start position plus the insertion length. We first recorded the observed number of overlaps between SVs and UTR, exon, intron, H3K4me3 and H3K27ac peak regions, respectively. Then we counted the average number of overlaps between randomly shuffled SVs (1,000 times) and each genomic region.

### SV genotyping in whole genome sequencing data

We collected whole-genome sequencing data for 643 domestic sheep, from four geographic regions (Europe, East Asia, Middle East and Africa) as well as wild sheep including 33 Asiatic mouflons (*Ovis aries musimon*) and 8 argali sheep (*Ovis ammon*). For the domestic sheep, we selected breeds with at least five individuals and defined populations by combining breeds from the same country with less than 5 individuals.

In this way, all the domestic sheep were assigned to 45 breeds or populations (Supplementary Data 7). We genotyped SVs in these samples using Paragraph v2.4a^32^. The maximum permitted read count for each variant was set to 20 times the average sample depth in order to reduce runtime for repetitive regions. The resulting genotype files in vcf format from all samples were then combined using bcftools v1.9^66^.

To obtain a high-confidence call set of SVs, the resulting genotypes were filtered based on overall genotyping rates and Hardy-Weinberg equilibrium^33^. Those variants 90% of all samples were retained using ≥ vcftools^67^ for further analysis. We also estimated Hardy-Weinberg equilibrium for each variant using vcftools and those with excess of heterozygotes (*P* value< 1×10^-5^) were removed.

### Quantifying linkage disequilibrium (LD) of SVs with surrounding SNPs

In order to measure the extent to which SVs are correlated with SNPs, we calculated LD (r^2^ values) between the polymorphic SVs (MAF>0.05) and the neighboring polymorphic SNPs (MAF>0.05; 150 upstream and 150 downstream SNPs) in Mouflon and domestic sheep. SNPs located on SVs were not considered. Plink v1.9 was used to calculate LD for all pairwise cases for SV-SNP and the surrounding SNP-SNP variant pairs. According to a previous study^41^, we first obtained pairwise genotype correlations (r^2^) for all SNP-SNP and SNP-SV pairs. For each of the 300 ranked surrounding positions, the number of times the SV rank was greater than the SNP-SNP median rank was recorded. Low, middle or high LD of each SV were then classified if the relative LD metric of SV to SNP was ≤ 100, > 100 and ≤ 200, and > 200, respectively, respectively.

### Inferring the derived SVs and *de novo* SVs in domestic sheep

To infer the state (ancestral or derived) for each SV, we genotyped one snow sheep (12.11× depth, ERR4161992) and a bighorn sheep (10.3× depth, SRR501898), which together with one argali sample served as outgroups. For each biallelic SV, if the same homozygous genotypes were detected in at least two outgroups, this state was defined as the ancestral state while the other was the derived state. If the derived allele is only present in domestic sheep but absent in all the wild sheep (8 Argali and 33 Asiatic mouflons), the SV was defined as a *de novo* SV in domestic sheep.

### Detecting selection signals

To detect population-stratified SVs, we employed two strategies, the global *F*_ST_ and the Ohana method, which identifies extreme allele frequency differences by modelling ancestral admixture components^36^. The global *F*_ST_ was calculated following a previously described approach with modifications^68^. For each SV site, we calculated the *F*_ST_ values between all populations. Instead of using the average of all *F*_ST_ values, we took the average of the top 50% *F*_ST_ values as the global *F*_ST_ value for each SV.

The top 1% of the highest global *F*_ST_ value by this method were considered as putative selective signals. For the Ohana method^36^, we first modeled the dataset with the provided workflow using the SNPs^33^. The resulting covariance was used as the neutral input for scans of selection. For each specific ancestry component, Ohana reported a likelihood ratio statistic (LRS) to quantify the likelihood of selection for each variant. We varied the number of ancestral admixture components from K=2 to K=5 to find selection signals.

In order to reveal selection signals in fat-tailed sheep, we used the method of population branch statistic (PBS) to calculate and compare the Fixation index (*F*_ST_) between the fat-tailed sheep, thin-tailed sheep and Asiatic mouflon, using the Asiatic mouflon as a distantly related population. The *F*_ST_ value between populations with different traits was calculated using the python scikit-allel package v1.3.2 (https://github.com/cggh/scikit-allel). We calculated the *F*_ST_ values of both SNP and SV sets. *F*_ST_ values for SNPs were calculated with a 10 kb sliding window with a 5 kb step.

### Detecting expression-associated SVs

The RNA-seq data of 8 tissues (liver, rumen, tail fat, skin, subcutaneous fat, semitendinosus, semimembranosus, longissimus dorsi) from 14 Texel (♂)×Kazak (♀) sheep hybrids with accession number PRJNA485657 as well as their whole genome sequencing data were downloaded from NCBI Sequence Read Archive^34, 43^. The raw reads were first trimmed for adapter and low-quality sequences using Trimmomatic^69^ and the high-quality reads were aligned to the sheep reference genome using Hisat2^70^. Reads counts per gene were computed using featureCounts ^71^ and normalized to counts per million (CPM) using edgeR package^72^ and those with CPM>1 in at least two samples were retained.

It is expected that the regulatory SVs could cause allelic-specific expression. We thus performed allelic specific expression (ASE) mapping by integrating the RNA-seq data as well as the SV genotyping information for each of the samples. The ASE genes were identified following the pipeline described in our previous study^34^. First, for each sample, the ASE SNPs displaying allelic imbalance at the gene body were identified from RNA-seq data using the following parameters: (1) at least 20 total reads; (2) allele ratio (allele reads count/total reads count)> 0.65 or <0.35; (3) significant allelic imbalances with FDR<0.05 (Chi-squared test). Those with at least two ASE SNPs at the gene body and found in at least three samples were defined as candidate ASE gene. For each SV-gene pair, if the ASE gene matches with the heterozygous state for the associated SV in at least five RNA-seq samples, we considered the SV as plausibly expression-associated.

The SV-based eQTL analysis was performed using the R package Matrix eQTL v.2.3 ^73^. The midpoint where SV starts and ends were defined as the position of each SV. Cis-eQTL analysis was tested within each tissue against all SVs located within the 200 kb upstream and downstream of neighboring genes. The gender and population structure (the first three principal components from PCA analysis of the 14 individuals) were included as covariates to account for confounding effects using an additive linear model. Statistical significance was considered with FDR<0.05.

### Analyzing the association between the 169 bp insertion with the tail length

The association analysis was conducted in a F2 population which was generated by backcrossing East Friesian sheep (♂) with the female Hu sheep (♀) × East Friesian sheep (♂). A total of 211 individuals of the similar age (six months old) were used. The tail length (cm) of each individual was measured and DNA samples were collected. Since the insertion was dominant in East Friesian sheep with allele frequency of 0.92 but is completely absent in Hu sheep based on our resequencing data, the F2 population that we studied only possessed two genotypes (II and ID) due to the grading-up using East Friesian sheep. Agarose electrophoresis was used to determine genotypes of the PCR products. Primers were designed in the flanking region of the insertion (forward: TTTATGAGCTTCTCTCCGCCA; reverse: AAGTGGTATAATTGCCGGGCT). PCR amplification was performed by denaturation of 94°C for 2 min followed by 28 cycles of 94°C for 30 s, 62°C for 30 s, and 72°C for 30 s and an extension of 72°C for 2 min; and a hold at 4°C.

### Luciferase reporter assay

To analyze promoter activity of the SV variation by luciferase assay, fragments with (mutation) and without (wild type) the insertion was synthetized by Tsingke Biotechnology Co., Ltd. (Beijing, China). Each fragment was cloned into the pGL3-Basic vector using Sac I and Hindu sites to assay its promoter activity on the downstream gene.

The 293T cells in 96-well plates were transiently transfected with 200 ng Firefly luciferase vector (pGL3) and 4 ng pRL-TK Renilla luciferase vector (Promega, Madison, USA) using 0.5 uL Lipofectamine 3000 (Thermo Fisher Scientific, Waltham, USA). Twenty-four hours after transfection, luciferase activities were measured with a dual-luciferase reporter assay kit (Promega, Madison, USA). Renilla luciferase activity was used as the transfection control for normalizing Firefly luciferase activity. The pGL3-Basic vector without insertion was used as the reference for promoter activity assays and the pGL3-Control vector was used as the positive control for promoter activity.

### Author contributions

Y.J. and S.Q.G. conceived and designed the experiments. R.L., X.M.Z. and Z.Y.L. performed *de novo* assembly and SV calling; Z.Y.L., M.S.X. and Y.F.Z. analyzed the RNA-seq and Chip-seq data; F.W., M.G. and L.Z. contributed to the ASE analysis; R.L., Z.Y.L., W.W.Fang and X.L.D. carried out SV genotyping and population genetics analysis; Y.T.Y. and X.Y.L. conducted the luciferase reporter assay; Z.B.Z., W.W.Fu, H.H.Z., C.N.C., P.Y., Z.A.G., N.J.N., H.A.N., X.P.Y., Y.X.S. and W.D.D. collected the samples and prepared the WGS data; R.L. wrote the manuscript and Y.J., S.Q.G., J.A.L., P.H., E.I.A., R.D.X. and X.H.W. revised it. All authors read and approved the final manuscript.

## Acknowledgements

We thank the High-Performance Computing platform of Northwest A&F University for providing computing resources. We would also like to thank Prof. Xiaolong Wang for providing the nice chart of different sheep tail patterns. This study was supported by research grants from the National Natural Science Foundation of China to Y.J. (U21A20247, 31822052), S.Q.G. (31760662, 31860639) and R.L. (31802027), Natural Science Basic Research Program of Shaanxi (K3030121801), and China Postdoctoral Science Foundation to W.W.Fu (No. 2021M702690).

## Ethics statement

Blood samples were taken by conforming with the Helsinki Declaration of 1975 (as revised in 2008) concerning Animal Rights.

## Data availability

Sequences and metadata generated in this work are publicly available. The *de novo* assemblies, HiFi data and WGS data for the 13 individuals are available at NCBI: WDU1 (PRJNA788705, PRJNA788704, PRJNA784371),QK1 (PRJNA788680, PRJNA788679, PRJNA784330), ROM1 (PRJNA788688, PRJNA788687, PRJNA784357), SFK1 (PRJNA788697, PRJNA788696, PRJNA784366), UJM1 (PRJNA788702, PRJNA788701, PRJNA784376), CHA1 (PRJNA788666, PRJNA788665, PRJNA784396), YUN1 (PRJNA788712, PRJNA788711, PRJNA784413), TEX1 (PRJNA788214, PRJNA788213, PRJNA780104), DOR1 (PRJNA788571, PRJNA788570, PRJNA782680), KER1 (PRJNA788676, PRJNA788675, PRJNA783516), MER1 (PRJNA788672, PRJNA788671, PRJNA783537), KAZ1 (PRJNA788564, PRJNA788563, PRJNA783179), FRI1 (PRJNA791176, PRJNA791175, PRJNA721520). The details are provided in Supplementary Table 1.

## Supplementary Files

**Supplementary Table 1** Accession numbers of the *de novo* assemblies.

### Supplementary Data

**Supplementary Data 1** Information of divergent variations.

**Supplementary Data 2** Information of multiallelic variations.

**Supplementary Data 3** List of the biallelic insertions and deletions identified in this study.

**Supplementary Data 4** SV frequency in 45 sheep breeds or populations.

**Supplementary Data 5** List of population-stratified SVs.

**Supplementary Data 6** List of putatively expression-associated SVs identified by ASE mapping.

**Supplementary data 7** List of putatively expression-associated SVs identified by eQTL mapping.

**Supplementary Data 8** Sample list of the resequencing data.

### Supplementary figures

**Supplementary Fig. 1** Example of three representative divergent variations. intersecting with IDE (a), PRP3-like (b) and MAGI2 (c), respectively.

**Supplementary Fig. 2** Example of three representative multiallelic variations affecting ASIP (a), UDP-glucuronosyltransferase 2B18-like (b) and olfactory receptor (LOC101115700) (c), respectively.

**Supplementary Fig. 3** The number of SVs intersecting functional elements compared to randomly permuting SV locations.

**Supplementary Fig. 4** An ideogram showing SV hotspots using all SVs. The total number of SVs in each detected hotspot is shown by a scale going from blue to red.

**Supplementary Fig. 5** Enrichment analysis of SVs with respect to the last 5 Mb of each chromosome end.

**Supplementary Fig. 6** PCA plot showing the clusters of domestic sheep by insertions (a) and deletions (b).

**Supplementary Fig. 7** Enriched KEGG pathways for the population stratified SVs.

**Supplementary Fig. 8** RNA-seq mapping for the 169 bp insertion shows that the insertion belongs to 5’ UTR region.

